# Two-Point Calibration Protocol for the FRET Indicator Pyronic in Neurons

**DOI:** 10.1101/2025.06.23.661049

**Authors:** F Baeza-Lehnert, Y Contreras-Baeza, C Aburto, A San Martín

**Author notes:** Corresponding authors: San Martín A.,; Baeza-Lehnert F.

## Abstract

**Significance:** Pyruvate is a nodal intermediate in cellular metabolism, positioned at the crossroads between glycolysis and fermentative metabolism. It is exchanged between the intracellular and extracellular compartments through the proton-coupled monocarboxylate transporters and between the cytosol and mitochondria through the mitochondrial pyruvate carrier, where it serves as a primary carbon source for respiration.

**Aim:** Our goal is to present a detailed protocol for quantifying cytosolic pyruvate concentration in neurons at single-cell resolution using a minimally invasive, two-point calibration approach with the FRET-based genetically-encoded fluorescent indicator Pyronic.

**Approach:** This protocol is based on a non-invasive pharmacological two-point calibration approach, where Pyronic’s dynamic range (ΔR_MAX_) is established by using trans-acceleration exchange to deplete intracellular pyruvate (R_MIN_), and by inducing Pyronic saturation (R_MAX_) through the combination of inhibition of pyruvate export, stimulation of its production, and blockade of its mitochondrial consumption. The protocol also incorporates the previously published K_D_ values for Pyronic obtained from *in vitro* experiments. This procedure does not require the use of detergents to permeabilize the cells.

**Results:** Implementing this protocol enables the measurement of absolute cytosolic pyruvate concentrations. This quantitative parameter facilitates comparisons of pyruvate metabolism across different cells, samples and experimental batches, thereby enabling the comparison between a plethora of experimental conditions.

**Conclusions:** The FRET-based fluorescent indicator Pyronic can be reliably calibrated using a minimally invasive, pharmacology-based two-point calibration protocol in neurons, thus providing a robust and quantitative method to study pyruvate metabolism under various physiological and pathological scenarios.

## 1 Introduction

Metabolism has recently gained renewed attention since metabolic dysfunction has been linked to the development of several pathologies, especially in the aging brain where glycolytic dysfunction has been detected [1, 2]. Pyruvate is a monocarboxylate critically located at the crossroads of glycolytic and fermentative metabolism. This monocarboxylate is a three-carbon metabolite produced in the cytosol of living cells by the breakdown of glucose, a six-carbon hexose through the glycolytic pathway. Through the lactate dehydrogenase (LDH) catalysis, it is reduced to lactate, a three-carbon metabolite. Depending on its concentration gradient across the plasma membrane, it can be exported to the extracellular space mediated by the proton-coupled monocarboxylate transporters (MCTs). Furthermore, driven by the mitochondrial pyruvate carrier (MPC), it feeds the mitochondrial tricarboxylic acid cycle (TCA), generating ATP or as a building block for cell growth. Therefore, pyruvate is a core metabolite at the intersection of glycolysis and oxidative phosphorylation (OXPHOS), displaying catabolic and anabolic functions. In oxidative cells, such as neurons, halting the provision of pyruvate into mitochondria leads to a rewiring of oxidative metabolism [3] and changes in synaptic transmission and neuronal excitability [4, 5] in both *in vitro* and *in vivo* models. Interestingly, pyruvate supplementation has been used in models of kindling and Alzheimer’s as a metabolic corrector [6–8].

Commonly, the fate of pyruvate metabolism has been traced using mass spectrometry, nuclear magnetic resonance (NMR), isotopic labelling, and colorimetric techniques. While, these approaches have proven useful, they present several limitations: low spatial resolution; requiring thousands of cells per data point, low temporal resolution; making it challenging to capture rapid events in the order of seconds, consume the sample; precluding before-and-after experimental designs, and require expensive equipment; preventing a broad accessibility to technology. Accordingly, fluorescent protein-based reporters have been engineered to measure pyruvate metabolism in single cells with second-scale temporal resolution, providing an easily detectable readout, with the potential to be well-calibrated without consuming the sample and the use of permeabilization procedures.

Genetically encoded fluorescent indicators are fusion proteins engineered from a binding-ligand domain and a fluorescent reporter module [9]. The first and most common architecture is the Forster Resonance Energy Transfer (FRET) [10]. In these types of indicators, the binding of the metabolite of interest induces a conformational change that alters the distance and orientation between the donor and acceptor fluorescent proteins, thereby affecting the FRET efficiency (**Figure 1A and B**).

**Figure 1.**
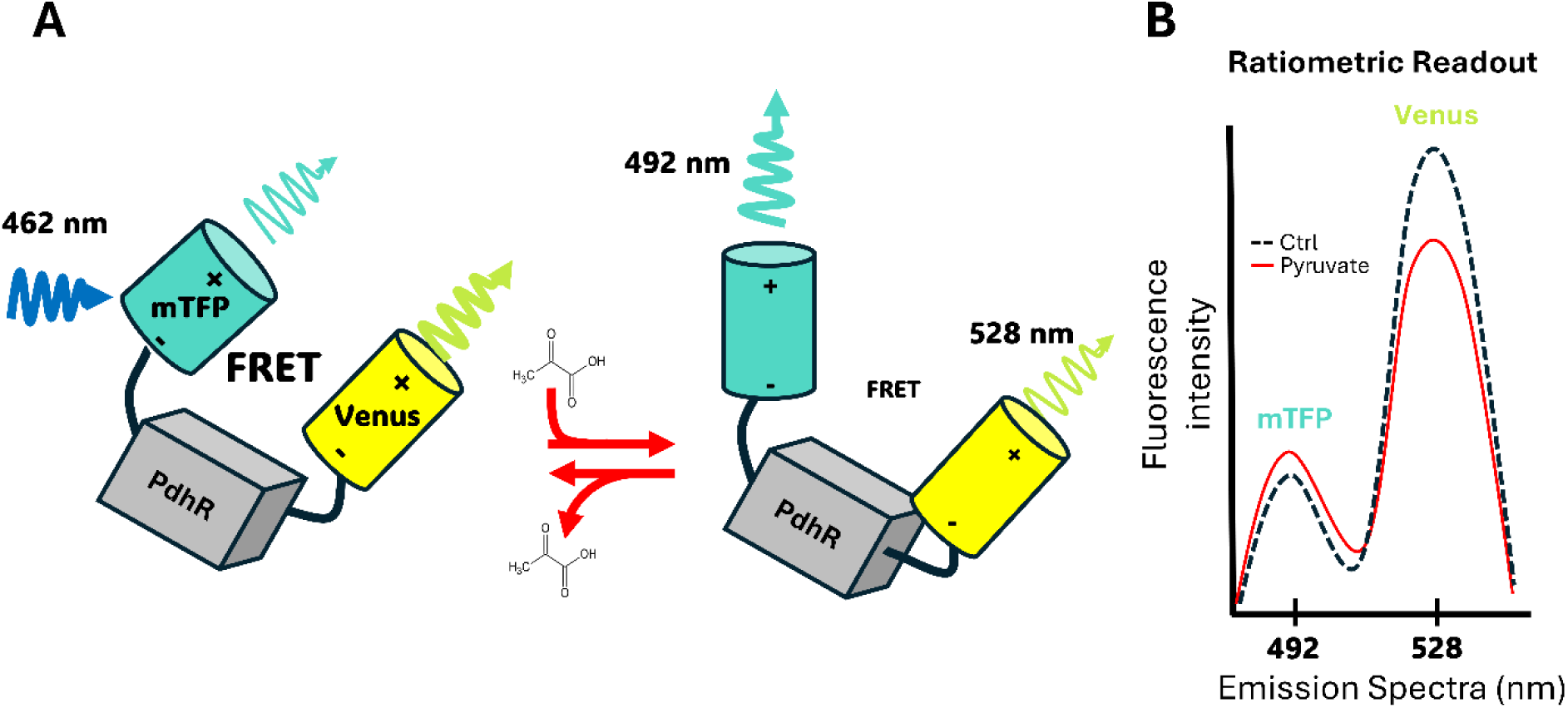
Schematic representation of the Pyronic and its spectroscopic properties. **A.** Pyronic consists of the metabolite-binding transcriptional regulator PdhR flanked by two fluorescent proteins, mTFP (donor) and Venus (acceptor). In the absence of ligand (left), the close proximity of the fluorophores enables efficient Förster resonance energy transfer (FRET): excitation at 462 nm predominantly excites mTFP, leading to strong Venus emission at 528 nm. Binding of pyruvate or lactate to PdhR (center) triggers a conformational change that increases the donor–acceptor distance and/or alters their relative orientation, thereby lowering FRET efficiency. This rearrangement increases mTFP emission at 492 nm and decreases Venus emission. **B.** Pyronic yields a ratiometric readout. The dashed black trace represents the basal (ligand-free) condition, whereas the red trace shows the fluorescence spectrum after pyruvate addition, demonstrating reciprocal changes in fluorescence intensity at 492 nm (mTFP) and 528 nm (Venus).

These changes in steady-state fluorescent intensity can be easily monitored using fluorescent microscopy, providing valuable readouts of relative levels, concentration, and fluxes. These fluorescent indicators have emerged as a convenient tool to assess rapid intracellular metabolite dynamics, and their readout is amenable to calibration for quantitative assessment of intracellular metabolites [9]. However, due to the complexity of cellular metabolism in terms of metabolite transport and metabolism, precisely controlling their intracellular concentration in a non-invasive manner is challenging.

The most popular readout from fluorescent indicators is fluorescent intensity, which is not inherently quantitative. However, using the same baseline signal it is possible to make quantitative comparisons in the same cell in a before-and-after experimental design [11–13]. Nonetheless, absolute numbers are needed because the behavior of the indicator is affected by the cellular microenvironment [14–16], protein folding and chromophore maturation [17, 18], equipment optics [19], and other factors [20], making it impossible to compare cells and sample batches directly using fluorescent intensity or FRET-ratio (**Figure 2**). Converting fluorescence data into metabolite concentrations permits wider comparisons between studies and experimental conditions, as well as quantitative analysis of metabolic networks. Therefore, quantitative assessment is highly desirable to avoid artefacts. If the intracellular level of a given metabolite can be manipulated—either depleted or loaded—a one-point calibration can be performed by measuring the apo- or saturated-indicator signal. In both cases, additional information, such as the dynamic range and K_D_ from the original characterization using a purified fluorescent indicator, is required. If it is possible to obtain both the apo- and saturated-indicator signal by manipulating the metabolite’s concentration from the extracellular space, a two-point calibration can be performed, relying on experimental data except for the pre-established K_D_. Additionally, full fluorescent indicator calibration is another alternative that involves detergents to permeabilize the plasma membrane and the superfusion of intracellular-emulating solutions, containing known metabolite concentrations, offered in a stepwise sequence. However, these procedures were found to be impractical and cumbersome due to rapid cellular swelling and sensor loss [21, 22]. In this regard, among the available experimental strategies for obtaining a quantitative assessment of a given intracellular metabolite, two-point calibration offers a good balance between invasiveness and the level of precision required for accurate evaluation of intermediate metabolism.

**Figure 2.**
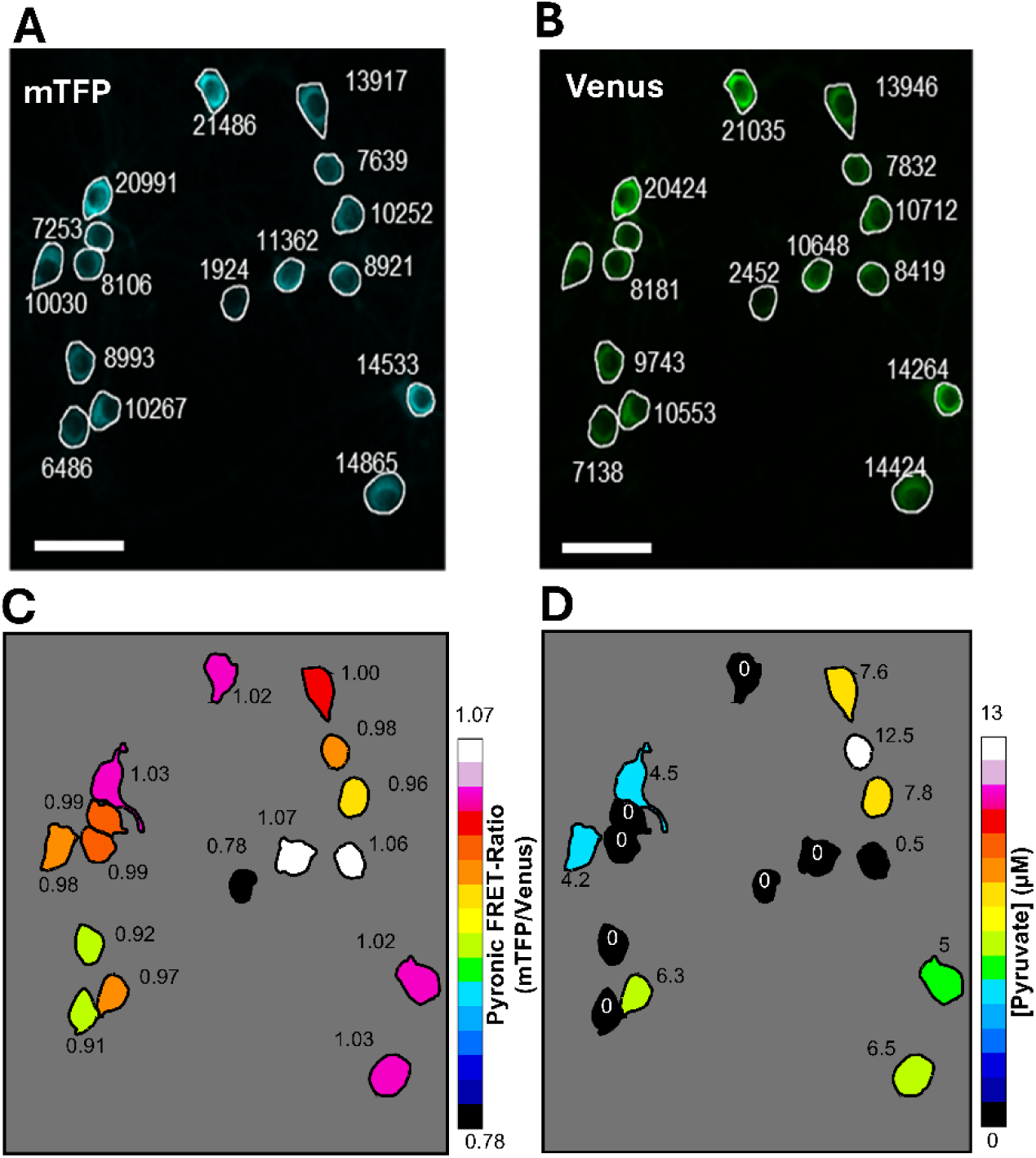
FRET ratio does not directly reflect intracellular pyruvate concentration. Confocal micrograph showing the cytosolic distribution of Pyronic in cultured neurons. Arbitrary unit of fluorescent intensity of **A.** mTFP channel **B.** Venus channel **C.** Pseudocolour representation of Pyronic FRET ratio. **D.** Pseudocolour map of the intracellular pyruvate concentration at single-cell resolution obtained using the two-point calibration protocol. The procedures involved to obtain absolute number will be disclose below. All the images were acquired under conditions of 2 mM glucose and 1 mM lactate. Scale Bar 30µm.

With the aim of monitoring and quantifying pyruvate dynamics with single-cell resolution, we engineered Pyronic [23], a pyruvate-sensitive genetically-encoded FRET-based sensor. This fluorescent indicator have been instrumental to study brain energy metabolism *in vivo* with cellular resolution [24–26], determine the affinity of monocarboxylate transporter [12], identification of a monocarboxylate transporter associated with type 2 diabetes [27], assessment of mitochondrial pyruvate consumption in neurons [28], energization endothelial cells motility [29], the influence of chronic exposure of ketones bodies in astrocytic mitochondrial metabolism [30] and the identification of neuronal signals of metabolic coupling [31, 32].

Here, we describe a minimally invasive protocol that allows for two-point calibration of Pyronic FRET signal from neurons. This protocol can be performed *in vitro* and *ex vivo* preparations and extended to other brain and non-brain cells.

## 2 Pyronic, a FRET-based fluorescent indicator for pyruvate

Pyronic is a genetically-encoded FRET-based sensor for pyruvate, based on the *Escherichia coli* transcriptional regulator PdhR, flanked with the fluorescent proteins mTFP and Venus, as FRET donor and acceptor fluorophores, respectively (**Figure 1A**). Pyruvate binding triggers a conformational change that lowers FRET efficiency, thereby increasing donor (mTFP) fluorescence and concomitantly decreasing acceptor (Venus) emission detected in the FRET channel (**Figure 1B**). Pyronic affinity constant (K_D_) for pyruvate of 107 ± 13 µm. Due to the dependency of the FRET ratio on pyruvate concentrations, Pyronic can detect pyruvate concentrations ranging from 10 to 1000 µm, which nicely covers the entire physiological range of cytosolic pyruvate in mammals and other systems, typically in the range of 20-200 µm [33]. Pyronic is highly selective, showing no sensitivity for other physiologically relevant monocarboxylates, and it is insensitive to pH within the physiological range [23]. Pyronic shows approximately 20% of maximum dynamic range (ΔR_MAX_) on purified protein and 40% in astrocytes, HEK293, COS7 and MDA-MB-231 cells [12, 23, 27, 33]. However, in neurons Pyronic showed a differential dynamic range between 20-35% [25, 28, 34]. The variation in dynamic range stems from the different expression levels reached with the selected gene-delivery methods. Transfection using cationic lipids typically introduces enough gene copies to yield high levels allowing it to reach a dynamic range of roughly 40 %, whereas viral vectors usually achieve only 20–35 %. Therefore, a two-point calibration is ideal for calibrating the fluorescent signal, because the dynamic range depends on the indicator expression level attained in each individual cell. Determining the dynamic range on a cell-to-cell basis prevents over- or underestimation of the fluorescence window used to accurately calculate intracellular pyruvate concentrations.

## 3 Rationale for the Two-Point Noninvasive Calibration of Pyronic

Cytosolic steady-state level of pyruvate is sustained by glycolytic production, the equilibrium through the LDH reaction with lactate, and the transport across the plasma and the inner mitochondrial membranes by the MCT2 and MPC transporters, respectively. We have devised a simple two-point calibration protocol based on this simplified metabolic arrangement to assess the minimum and maximum Pyronic signal.

Given the high-affinity MCT2 transporter and the pyruvate oxidation within mitochondria, cytosolic pyruvate quickly drops, and Pyronic fully desaturates when neurons are superfused with a saline solution without energy sources, allowing the R_MIN_ (mTFP/Venus minimal ratio, apo-Pyronic state) to be obtained. On the other hand, when MCT and MPC are pharmacologically blocked while glycolysis is boosted to increase pyruvate production, cytosolic pyruvate concentration saturates Pyronic to obtain the R_MAX_ (mTFP/Venus maximal ratio, fully saturated-Pyronic state). To afford this, we bath neurons with a saline solution containing 10 mM pyruvate, 5 mM glucose, 1 µM AR-C155858 or AZD3965 both MCT1/2 blockers [35, 36], 10 µM UK-5099 an MPC blocker [37], and 5 mM sodium azide. Sodium azide is a mitochondrial complex IV inhibitor that stops the electron transport chain (ETC), halting mitochondrial ATP production. However, due to the drop in cytosolic ATP, glycolysis is boosted as a metabolic compensation [38], favoring pyruvate production and therefore its intracellular accumulation. All together, these procedures performed at the end of each experimental run revealed the dynamic range of Pyronic ΔR_MAX_ (R_MAX_ - R_MIN_), uncovering its full dynamic range in each cell from every batch of samples. Determining the full dynamic range of Pyronic from each cell is important, especially when a heterogeneous fluorescent signal is detected depending on the gene-delivery system. Although this design was developed for primary hippocampal neurons, it could be applied to any other cell type by simply adjusting the MCT inhibitor to target the appropriate isoform.

## 4 Primary hippocampal neurons from mice

Primary cultures offer a favorable balance between experimental control and physiological relevance. Consequently, this model allows direct and controlled access to the cellular environment, enabling the establishment of causality for a given pharmacological, nutritional, optogenetic, or genetic perturbation. This makes it an ideal model to elucidate the nuances of the mechanisms underlying a phenomenon of interest. Additionally, we have gravitated to co-culture embryonic hippocampal neurons and astrocytes to promote *in vitro* differentiation and metabolic cooperation [28, 39, 40]. This type of co-culture of neurons and astrocytes has been used to unveil a plethora of neuronal signals related to metabolic coupling, which have been validated *in vivo* [25, 41].

A detailed description of the steps involved in sample preparation for the imaging experiment will be provided below.

### 4.1 Experimental Model and Subject Details

Hybrid females F1 C57BL/6J x CBA/J mice aged 2-6 months carrying 6-9 embryos were used. All animals were housed in standard pathogen-free (SPF) conditions with a 12:12-hour light/dark cycle at room temperature (20 ± 2°C) and had free access to food and water. All experimental procedures were approved by the Centro de Estudios Científicos Animal Care and Use Committee following the recommendations of the Guide for the Care and Use of Laboratory Animals, Institute of Laboratory Animal Resources, National Research Council.

## 5. Drugs, reagents and equipment

### 5.1 Euthanasia

- Isoflurane (100%, UPS grade, Baxter cat. #10019036060)
- Anesthesia chamber
- Cotton pads or gauze
- Pasteur pipette
- Cervical Dislocation Support Base
- Dedicated protective clothing for the dissection area (e.g., face mask, gloves, lab coat)
- Disposal container for carcasses

### 5.2 Dissection

- Dissecting microscope (e.g., Olympus SZ61)
- Large, sharp scissors
- Dissection pins
- Rat-tooth forceps (toothed forceps)
- Curved forceps and fine-tip forceps
- Fine scissors
- Two Petri glass dishes (100mm)
- Two 15 mL conical tubes, filled with cold Hanks’ Balanced Salt Solution (HBSS)
- Two 50 mL Falcon tubes, filled with cold HBSS.
- A Styrofoam box filled with ice
- Laminar flow cabinet

### 5.3 Neuronal culture

- 30 mm petri dishes (or alternatively, 6-well plates).
- 15 mm glass coverslips
- Poli-L-lysine (Sigma, cat. # P4832)
- Trypsin-EDTA (0.5%) in DPBS (10X) (Capricorn, Cat. TRY-1B10)
- Neurobasal-A medium, no D-glucose, no sodium pyruvate (Gibco, cat. # A2477501)
- D-Glucose solution (Gibco, cat. # A2494001)
- Sodium pyruvate (Gibco, cat. # 11360070)
- GlutaMAX Supplement (Gibco, cat. # 35050061)
- B-27 Supplement (Gibco, cat. # 17504044)
- Penicillin/Streptomycin (Gibco, cat. # 15140122)

### 5.4 Neuronal transduction with Pyronic

- Pyronic (Addgene: Plasmid #51308)
- AAV-DJ-hSyn-Pyronic
- Hippocampal mix-cultures

### 5.5 Saline solutions

- ddH_2_O
- NaCl (Sigma aldrich, CAS# 7647-14-5)
- KCl (Sigma aldrich, CAS# 7447-40-7)
- CaCl_2_ (Merck, CAS# 100-04-8)
- MgCl_2_ (Merck, CAS# 7791-18-6)
- HEPES (Merck, CAS# 7365-45-9)
- NaHCO_3_ (Merck, CAS# 144-55-8)
- NaOH (Merck, CAS# 1310-73-2) to titrate the saline solution to the corresponding pH.

**Caution**: NaOH is a strong base. We recommend using a 1M stock solution. Handle with care and wear appropriate PPE, including gloves, a lab coat, and eye protection.

- pH meter (e.g., Extech Model 321990)
- Osmometer (e.g., Advanced instrument Model 3250)
- Plate stirrer
- Analytical balance (laboratory scale)
- 1 L glass bottles
- beakers, volumetric flasks, spoons and spatulas, magnetic stir, plastic plates for measuring compounds, etc.

### 5.6 Drugs

- Sodium azide (Sigma aldrich, CAS # 26628-22-8)
- UK-5099 (MedChemExpress, CAS # 56396-35-1)
- AR-C155858 (MedChemExpress, CAS # 496791-37-8)
- AZD 3965 (MedChemExpress, CAS # 1448671-31-5)
- Dimethyl sulfoxide (Sigma-Aldrich, CAS # D8418)

### 5.7 Imaging Setup

- Wide-field epifluorescent microscope (e.g., Olympus BX51)
- Light Source (Xenon arc lamp)
- Excitation filter (Chroma Technology Corp. Scan range: 400 nm to 600 nm. Set: 51017bs)
- Dichroic mirror
- Emission filters (Chroma Technology Corp. Scan range: 400 nm to 600 nm. Set: 51017m)
- CCD camera (Rolera MGi-Plus EMCCD) Alternatively,
- Laser-scanning microscope (e.g., Fluoview FV1000)
- 440 nm solid-state laser (405 nm solid-state laser can be used instead; however, only with 50% efficiency in excitation)
- Excitation-emission cube: 405-440/515 dichroic mirror
- Emission split: SDM510 dichroic mirror
- Band-pass filters; mTFP: 465-495 nm, Venus: 535-565 nm.

### 5.8 Recording chamber and perfusion system

- Recording chamber (Warner instruments, RC-26GLP cat. #64-0236)
- Bipolar Temperature Controller Model CL-100 (Warner Instrument, cat. #W464-0352)
- Liquid Cooling System Model LCS-1 (Warner Instrument, cat. #W464-1922)
- Dual In-line Solution Heater/Cooler Model SC-20 (Warner Instrument, cat. #W464-0353)
- Thermistor Model TA-29 (Warner Instrument, cat. # W464-0107)
- Peristaltic and/or vacuum pump (Fisherbrand™ Variable-Flow Peristaltic Pumps). Alternatively, gravity-driven inflow can be used.
- 5% CO_2_/95% air gas mixture

### 5.9 Software

- Image acquisition and analysis software (dedicated software provided by the microscope or µManager ImageJ).

## 6 Equipment setup

### 6.1 Imaging setup

The imaging setup must include all the previously detailed optics to effectively record Pyronićs signal. Alternative setup arrangements, such as those based on LED illumination, optoscan monochromators and optosplits, are also suitable and amenable to FRET recordings. A quick check out of all the electronic systems, such as shutters and camera idling, as well as excitation and collection of the emitted photons, is highly recommended in an early introductory session.

On the day of recording, it is recommended to turn on all the equipment, including the microscope, light source, temperature regulator, and computer. Typically, it takes about 30 minutes for the microscope’s body temperature and light source power to stabilize. To minimize focus drift during experiments, an autofocus system can be attached to the microscope. The room should be darkened, and the temperature should be maintained at a stable level; using an air conditioning system is advisable.

### 6.2 Recording chamber

Disassemble the recording chamber and add vacuum grease to the chamber slit to secure a good seal. Inflow and outflow perfusion lines must be connected to the corresponding peristaltic or vacuum pumps and tested for proper functioning. The solution overflow regularly occurs because of poor arrangement of the tubing in the pump slits or connections.

### 6.3 Perfusion system

We use an open perfusion system that collects the working solution at one end, bathes the recording chamber, and then discards the solution at the other end (**Figure 3**). Another feature of our perfusion system is that the recording chamber is fully accessible with a pipette, which is beneficial for incubating drugs directly within the chamber when continuous perfusion is unnecessary or should be avoided. This situation may arise when testing a small amount of a drug or during extended preincubation times. However, it’s important to note that stopping the perfusion system when using bicarbonate-buffered solution can affect the pH. Therefore, we recommend using a HEPES-buffered solution in these cases. Alternatively, recirculation systems can be employed with bicarbonate-buffered solutions when dealing with small drug quantities or prolonged incubation times cannot be avoided. In this setting, both ends of the perfusion system must be placed in the same cylinder tube.

**Figure 3.**
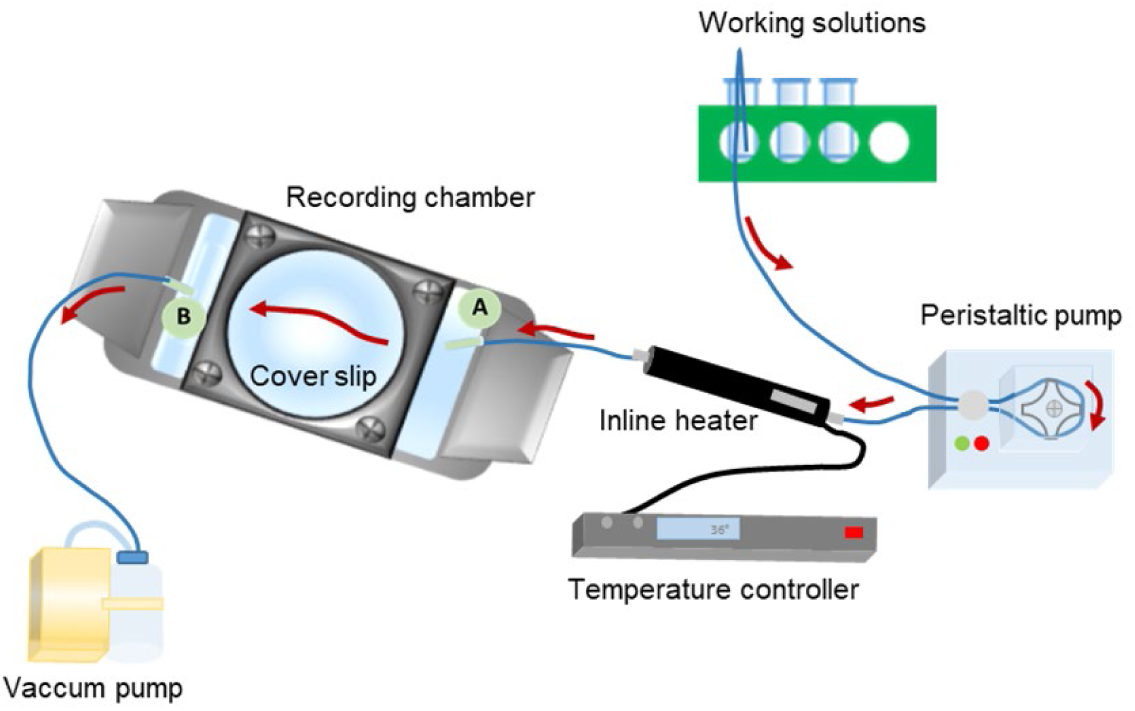
Open perfusion setup for single-cell recordings. Working solution is drawn from a reservoir through 1.6 mm inner-diameter silicone tubing and driven by a peristaltic pump toward the recording chamber inlet. When temperature control is required, the line passes through an inline heater placed as close as possible to the chamber. The solution bathes the fully accessible recording chamber, allowing localized drug application with a pipette when continuous flow is unnecessary. Effluent exits the chamber via an outflow line to a waste container or vacuum trap. Red arrows indicate the direction of the buffer flow.

Typically, the perfusion is established using 1.6 mm diameter silicone tubing. The first segment (hereafter, the inflow) goes from the cylinder tube containing the working solution, passes through the peristaltic pump, and ends at the inlet of the recording chamber. If temperature needs to be controlled, an inline heater should be placed between the peristaltic pump and the recording chamber. In this case, the inflow tubing is connected to the inline heater after being wrapped through the peristaltic pump slots. We recommend strategically placing the inline heater close to the recording chamber. Finally, a short piece of silicon tubing is needed to connect the inline heater to the inlet port of the recording chamber. The second segment, hereafter referred to as the outflow, extends from the outlet port of the chamber to the vacuum pump port or the disposal container.

### 6.4 Solutions

We commonly use bicarbonate buffer of the following composition (in mM): 112 NaCl, 3 KCl, 1.25 CaCl_2_, 1.25 MgSO_4_, 10 HEPES, 24 NaHCO_3_, pH 7.2 adjusted with 1M NaOH, and bubbled with 95% air / 5% CO_2_. Alternatively, a bicarbonate-free solution of the following composition can also be used (in mM): 136 NaCl, 3 KCl, 1.25 CaCl_2_, 1.25 MgCl_2_, 10 HEPES, pH 7.4 adjusted with 1M NaOH. We typically prepare 2 L of saline solution, which, depending on the experimental flow rate of the perfusion system, lasts approximately 15 hours of recordings. Importantly, we do not recommend using saline solutions older than two weeks, as the pH slowly changes, accompanied by the concomitant formation of a precipitate.

On the experimental day, fill a 100 mL volumetric flask with the saline solution of choice (bicarbonate- or bicarbonate-free-buffered) and supplement it with 2 mM glucose and 1 mM lactate. Next, split it into two 50 mL conical tubes, and add 10 mM pyruvate, 2 µM AR-C155858 or AZD 3965, 10 µM UK-5099 and 5 mM sodium azide into one of the two tubes. Finally, fill a third 50 mL conical tube with the saline solution of choice, but do not supplement with energy substrates. Once ready, all solutions must be placed in a rack by the imaging setup. Bicarbonate-buffered solutions must be bubbled with 5% CO_2_/95% air for at least 30 min to stabilize pH.

**Tip:** The experimental solution can be prepared after turning on the equipment to use the time more efficiently.

**Tip:** We use prepared 1 M stock solutions stored at -20°C for up to 6 months for glucose, lactate and pyruvate. Drugs are prepared as 1000x stocks and kept at -20°C.

## 7 Procedure

### 7.1 Mixed neuron-astrocyte primary cultures. (14-16 days before experimental day)

1. Pregnant mice at embryonic day 17.5 (E17.5) were euthanized by cervical dislocation following deep anesthesia with isoflurane, in accordance with institutional ethical guidelines and the ARRIVE principles. Under sterile conditions, 6–8 embryos were harvested and transferred to ice-cold Hanks’ Balanced Salt Solution (HBSS) supplemented with 5 mM glucose. Brains were rapidly isolated, and hippocampi were dissected under a stereomicroscope, ensuring complete removal of the meninges. Tissues were enzymatically dissociated in 1% trypsin-EDTA (Sigma-Aldrich) in HBSS for 15 minutes at 37°C. The enzymatic reaction was quenched by adding Neurobasal medium (Gibco) supplemented with 10 mM glucose, 2% B-27 supplement, 1% GlutaMAX, and 5% fetal bovine serum (FBS). Hippocampal cells were mechanically triturated and seeded onto 25 mm poly-D-lysine-coated glass coverslips at a density of 160,000 cells/mL. Cells were allowed to adhere for 2 hours at 37°C in a humidified incubator (95% air, 5% CO₂). After this period, the plating medium was replaced with serum-free Neurobasal medium supplemented with 10 mM glucose, 2% B-27 supplement, 1% GlutaMAX, 2.5 μg/mL and 10 μg/mL penicillin/streptomycin. Neuron-astrocyte co-cultures were maintained at 37°C in a humidified incubator (95% air, 5% CO₂), with two-thirds of the medium exchanged every 3 days.

### 7.2 Pyronic transduction in primary hippocampal neurons. (5-7 days before experimental day)

2. Add the AAV-DJ-hSyn-Pyronic particles directly to the culture medium 7 to 5 days before the experimental session and mix thoroughly. This procedure results in more than 80% of neurons expressing the sensor. Alternatively, lentiviral vectors or conventional lipofection procedures have been successfully used [23, 28]. **Tip:** The number of viral particles will depend on the titter of the viral preparation. We recommend using the lowest amount of virus that allows for good expression and a reasonable signal-to-noise ratio, without any observable inclusion bodies or signs of saturation. **Tip:** AAV serotype 2 is commonly used in cultured brain slices and stereotactic procedures due to its broad tissue tropism, enabling the efficient transduction of neurons. However, its transduction efficiency in primary neuronal cultures is limited. To overcome this, we cloned Pyronic into an AAV vector expressing the AAV-DJ capsid, a synthetic hybrid serotype generated by DNA family shuffling of eight wild-type AAVs (AAV1–9). AAV-DJ combines the advantageous properties of these parental serotypes to achieve high transduction efficiency across a broad range of cell types, including neurons. It exhibits superior *in vitro* infectivity compared to natural serotypes, making it an ideal vector for delivering genes to neurons in both dissociated cultures and tissue slices [42]. **Note:** In our hands, lipid transfection (Lipofectamine) is also possible, but the efficiency is low (<10%). **Tip:** The gene-delivery system determines the number of copies of the DNA delivered into the cell and, consequently, is one factor that determines the levels of expressed protein and the signal-to-noise ratio. Lipofection displays low transfection efficiency but allows obtaining heterogeneous and high fluorescent signals in some cells, resulting in ΔR_MAX_ values closer to the 40% dynamic range previously reported in various cellular systems. On the other hand, lentiviral particles offer high gene-delivery efficiency but yield a lower signal-to-noise ratio due to the limited gene copies that can be inserted into the genome, with a ΔR_MAX_ of approximately 20%. In our experience, adeno-associated viruses performed optimally, effectively balancing gene-delivery efficiency and signal-to-noise ratio, achieving a ΔR_MAX_ of around 35%. **Tip:** Cytosolic pyruvate concentrations have been reported in the micromolar range [28, 33]. Therefore, performing experiments in cells that display fluorescent indicators expressed in the same range could produce buffering effects [43]. This is the case for plasmids with strong promotes such as CMV or CAG that produces a micromolar range of fluorescent indicators [44, 45]. Under these conditions Pyronic could act as a chelator, decreasing the baseline levels of pyruvate and potentially affecting cell physiology. Additionally, cells overexpressing fluorescent indicators also exhibited significant impairment in mitochondrial oxidative respiration and changes in proteomics profile, suggesting broader mitochondrial dysfunction [46]. To minimize these scenarios, the use of gene-delivery systems such as adeno-associated virus with expression driven with cell-type specific promoter will minimize these unwanted effects.

### 7.3 Imaging Pyronic (On the experimental day)

3. Take the 25 mm glass coverslip from the petri dish and mount it in the recording chamber. Once the chamber is closed, add 0.5 mL of saline solution supplemented with 2 mM glucose and 1 mM lactate to prevent the cells from drying out.
4. Place the recording chamber in the microscope stage and connect the inflow and outflow lines. Inline heaters are typically positioned before the recording chamber if temperature control is desired. A thermistor is also recommended in the recording chamber to monitor temperature variations throughout the experiment. **Tip:** Experiments performed at 35–36°C are often noisier and more technically challenging. As temperature increases, the metabolic rate generally accelerates due to enhanced enzymatic activity, membrane fluidity, and faster molecular diffusion [47]. We recommend running a few experiments at room temperature when inexperienced with live imaging. This favors more stable recordings and improves the degree of confidence with handling the technique. To avoid degassing, heat the solutions in a water bath.
5. Turn on the perfusion system. The volume of superfusate in the chamber is approximately. 0.5 mL. Keep a stable and continuous perfusion rate of about 2 mL/min, which secures an approximate turnover time of the fluid in the chamber of about 15 s. Quick wash-in of sugars and drugs is desired for kinetic analysis of pyruvate dynamics; therefore, turnover times must be maintained high. **Tip:** Cells can also be superfused by gravity. Height has to be arranged accordingly to ensure a constant flux in the chamber, which must be correctly equalized to the efflux rate. Also, the speed of gravity-based perfusion decreases when saline solutions are used. The variation in the speed may induce substantial changes in temperature when an in-line heater is positioned before the recording chamber. **Tip:** When culturing primary brain cells, we typically use glucose concentration of 10 mM, but experiments are conducted in 2 mM glucose. Therefore, it is recommended to allow the cells equilibrate to the new conditions in about ten minutes. Notably, we observed that when fluorescent recordings of glucose began immediately after mounting the coverslip in the recording chamber, there was a gradual transition in intracellular glucose concentration towards a new steady state. This transition occurs as the cells need to reestablish glucose gradients. This issue may be even more pronounced when neurons are cultured in standard 25 mM glucose medium but are recorded at lower glucose concentrations.
6. Find the focal plane under bright field illumination. Using a 20 x objective is sufficient to record from neuronal bodies in primary culture. This also secures about 20 cells per field of view, improving data collection per experiment. The 40x objective is equally suited, but fewer cells are recorded per imaging session.
7. Switch on a light source (Arc xenon lamp or laser line), excite the sample at 440 nm and scan the plate using either of the two emission channels (mTFP or Venus). Alternatively, the sample can be scanned first with 480 nm excitation light and the Venus filter. Under the fluorescent microscope, Pyronic expression should be observed as homogeneously distributed in the cytosol and with explicit nuclear exclusion (**Figure 2A and B**). Bright spots indicate protein aggregation presumably due to overexpression, and should be avoided. **Tip:** When Pyronic is allowed to express for too many days (in our experience, more than 12 - 14 days), the sensor coalesces into large pyruvate-insensitive aggregates. This negatively impacts the sensor’s dynamic range and decreases the signal-to-noise ratio. We do not recommend imaging these neurons.
8. Take a snap with the imaging software and check for the level of fluorescence in each channel. Modify the illumination intensity or detector sensitivity accordingly to improve photon collection for both emission channels, preventing saturated pixels. As a general rule, illumination should be kept to the minimum amount of light that generates a reasonable signal-to-noise ratio. Excessive exposure can lead to photobleaching and/or phototoxicity, as well as contamination by autofluorescence, so it should be prevented. **Tip:** The two fluorescent proteins used in Pyronic are resistant to photobleaching; however, low illumination exposure times and power is always advisable. When illumination has to be strong, lower the acquisition frequency to minimize bleaching. Most commonly, intense illumination is needed when the Pyronic expression level is low; therefore, the transduction process may need to be reassessed.
9. Adjust imaging frequency according to the expected kinetics of the question at hand. Most commonly, frames every 5 to 10 seconds are sufficiently fast to follow the metabolic response. Images are taken regularly as 512 x 512 pixels, which helps monitor cell shape and movement throughout the experiment without storing unnecessary details. **Tip:** Depending on the signal-to-noise ratio, binning might be advisable. **Tip:** Most dedicated imaging software permits monitoring the experiment by a live plot of the Pyronic ratio (mTFP/Venus). This allows for real-time assessment of pyruvate dynamics, which eases the recognition of steady states or the effect of incubation with different saline solutions.
10. Start the imaging and wait for approximately 5-10 minutes, or until a stable baseline is reached. Ensure that the perfusion system runs steadily and that the solution volume trapped in the chamber and the temperature are constant during the experiment.

### 7.4 Pyronic two-point calibration

11. From a suitable field with non-saturated and non-inclusion bodies neurons expressing Pyronic (**Figure 2A and B**) get a stable baseline and switch to the solution without extracellular carbon sources for about 5 to 10 minutes to reach the minimum Pyronic signal (R_MIN_ - apo-Pyronic state). **Tip:** The time may vary from preparation to preparation due to the degree of spontaneous activity in the cultures. This step may take even longer if the experiments are performed in a priori silenced neurons (e.g., TTX treatment).
12. Next, switch to a saline solution containing 5 mM sodium azide, 1 µM AR-C155858 or AZD 3965 and 10 µM UK-5099, in the presence of 2 mM glucose, 1 mM lactate and 10 mM pyruvate, for 5 minutes to reach the maximum Pyronic signal (R_MAX_ - saturated-Pyronic state) (**Figure 4A**). As one example, when intracellular pyruvate precursors and inhibitors are offered sequentially, the increment of Pyronic signal, from minimum to maximum, can be unveiled as shown in **Fig. 4B**. The same maximum signal is obtained when the pharmacological calibration cocktail is administered together. Thus, we recommend this simplified single-step procedure when assessing the maximum Pyronic signal.
13. **Important note:** Assessing the maximum Pyronic signal is a final procedure, as cells must be incubated with a cocktail of drugs that bind with a high degree of affinity to their targets and presumably irreversibly modify metabolism. We strongly emphasize that the maximum point calibration procedure must be performed at the end of the experiment. Interpretation of the data afterwards will be dubious otherwise. **Note:** Pyronic does not exhibit proton sensitivity, which is important because certain protocol steps—such as 5 mM azide pulse—can produce acute intracellular pH changes. Therefore, the presented protocol is not advisable to be performed using pH-sensitive fluorescent indicators for pyruvate [33, 48–50].
14. Finalize the experiment and save the images for later analysis. If a live plot is unavailable, we recommend an on-the-fly analysis of the experiment before pursuing another one. Most dedicated imaging software is implemented with an analysis section where plots of the single channels can be evaluated. This quick visualization of the experiment allows for checking the correct responsiveness of the sensor and considering changes to the protocol, such as timing or illumination settings. **Important note:** We commonly perform the two-point calibration as a sequential two-step protocol at the end of the experiment. Nevertheless, depending on the question in mind, an ad-hoc protocol must be designed. As stated in section 3, this calibration procedure relies on the cellular response. Therefore, both the minimum and maximum Pyronic signals must be assessed when the conditions are most favorable, but without interfering with a clear-cut assessment of the biological question at hand. For instance, if the effect of a mitochondrial blocker is to be assessed, the minimal Pyronic signal must be evaluated before inhibiting mitochondria.

**Figure 4.**
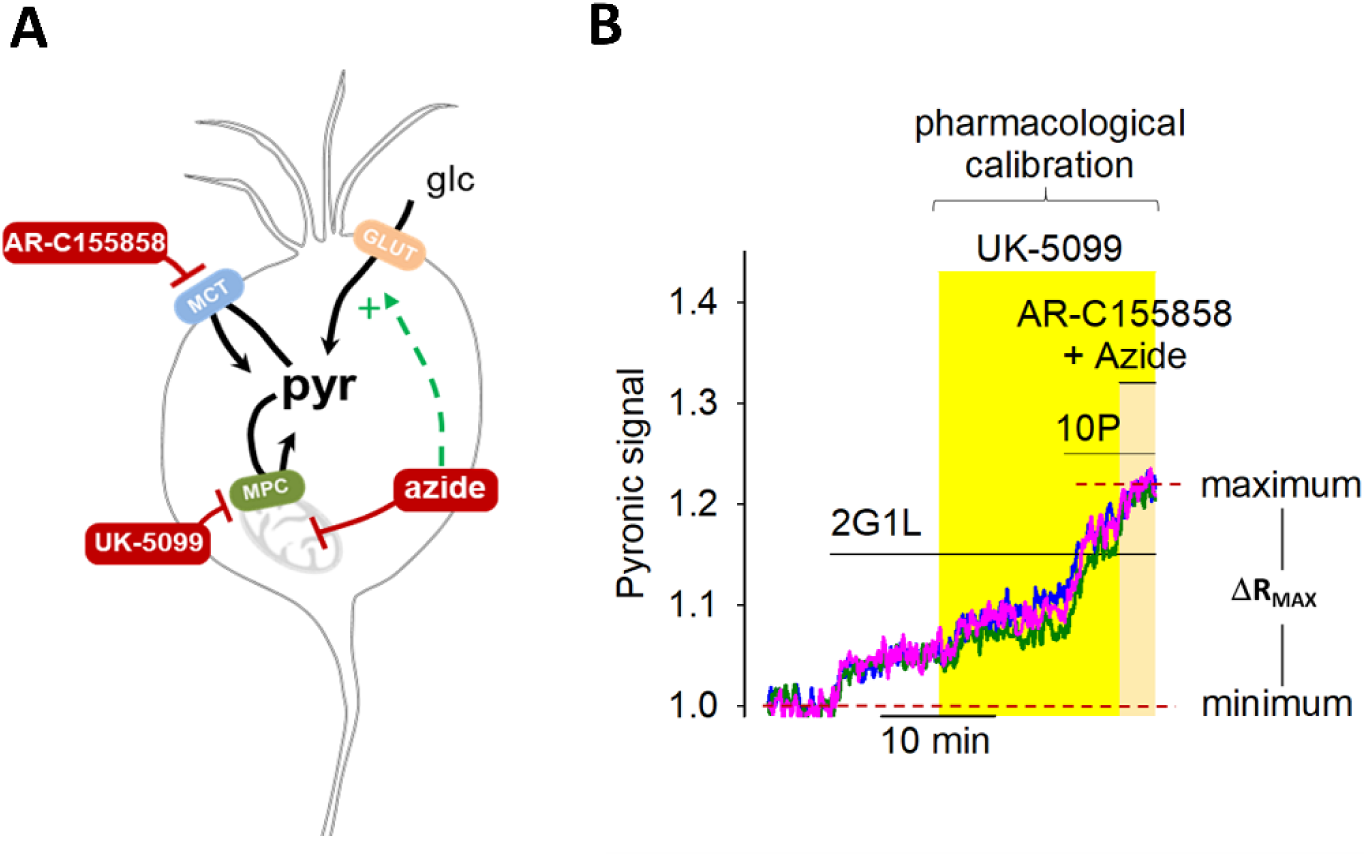
Pyronic two-calibration protocol. **A.** Schematic representation of the pharmacological protocol used to accumulate intracellular pyruvate. Glucose (glc) and lactate enter the neuron via GLUT and MCT transporters, respectively. Pyruvate is transported into mitochondria by the MPC. Pharmacological agents used include 1 uM AR-C155858 (MCT inhibitor), 10 uM UK-5099 (MPC inhibitor), and 5 mM azide (OXPHOS inhibitor). **B.** Representative traces of Pyronic signal from three single-neurons during the calibration protocol. The R_MIN_ (minimum) is established by removing extracellular glucose and lactate (2G1L condition). The R_MAX_ (maximum) is then achieved by sequential application of 1 µM UK-5099, 10 mM pyruvate (10P), 1 µM AR-C155858, and 5 mM Azide in the presence of 2 mM glucose and 1 mM lactate. The dynamic range is defined by ΔR_MAX_ = R_MAX_ - R_MIN_.

### 7.5 Data analysis: regions of interest and fluorescent intensities analysis

15. Open the file of images in an imaging analysis software or FIJI [51]. Depending on the software setting, the image stack will be automatically split into two channels. If this is not the case, separate them accordingly.
16. Scroll back and forth through the image stack and look for changes in the X-Y axis, cellular volume and shape, and background fluorescence. Sometimes, small pieces of cultured tissue become detached from the glass coverslip and eventually cross the field of view. **Tip:** Drift in the X-Y axis can be easily corrected with image stack alignment plug-ins available for FIJI. **Tip:** Z-axis drift is more challenging to correct. The temperature stability of the saline solutions and the room play an essential role in the stability of the Z axis. Alternatively, autofocus systems are commercially available for most imaging setups.
17. Select regions of interest (ROI), including one ROI in an area without cells (**Figure 5**). Commonly, we draw ROIs that are slightly larger than the cells to keep them within the ROI when a slight movement is presented. Scroll back and forth through the image stack, checking that cells remain within the ROI throughout the experiment. **Note:** Pyronic displayed a homogenous cytosolic pattern with a clear nuclear exclusion. The ratio of nucleus:cytoplasm in neurons is high. Therefore, we prefer to cover the whole cell with the ROI in order not to lose any detectable signal from cytoplasm.
18. Compute the average pixel intensity for each ROI on both channels. We do not recommend computing the maximum value or the sum of pixel intensities for each region, since it weights outlier pixels.
19. Copy and paste the fluorescence intensity values into an Excel sheet for further analysis. Firstly, a background-corrected value is calculated by subtracting the background intensity from each channel in each region of interest (ROI) over time. Next, to obtain the normalized Pyronic signal, the complete experiment is divided by the average of the Pyronic signal at the minimum ratio (R/R_MIN_). **Tip:** Typically, we plot the background and the background-corrected mTFP and Venus fluorescence. The background should be constant unless the ROI is placed in an area where thin neurites are present, movement, or something crosses the field of view. For the mTFP and Venus fluorescence, changes in the intensity must be reciprocal between the channels, as a result of FRET. Pyronic is ratiometric, thereby insensitive to changes in cellular volume, drift and movement. **Tip:** Non-reciprocal changes must be evaluated carefully, specifically because of potential pH artefacts. Within the physiological range, Pyronic is insensitive to pH; however, substantial pH changes can affect the single fluorophores to varying degrees. The pH artefact manifests as quickly developing changes in the fluorescence of the two channels, typically in the same direction; however, the Venus channel is usually more affected. On occasions, experimental manipulation can induce a real change in the metabolic response, co-existing with a pH change; careful analysis and interpretation are advised until pH is controlled, or ruled out. Genetically encoded pH sensors and chemical dyes are available elsewhere, and controls of pH are highly recommended. **Note:** Pyronic is based on mTFP [52] and Venus [53]. FRET-pair. This fluorescent protein pair virtually eliminates signal contamination (“bleed-through”) for two spectral reasons. First, excitation at 458 nm coincides with the absorption maximum of mTFP (462 nm) but barely excites Venus, so the contribution of acceptor spectral bleed-through to the FRET channel is negligible. Second, although the mTFP emission peak is shifted 17 nm toward the green relative to CFP, its emission band is much narrower, meaning that overlap with the Venus detection window (≈ 535– 590 nm) is minimal [54]; this drastically reduces donor spectral bleed-through in the FRET channel. Therefore, spectral bleed-through correction is not necessary for Pyronic. In our setup, the mTFP was excited with a 440 nm solid-state laser, which did not induce excitation of Venus, and FRET emission was collected in a 535–565 nm window that overlaps minimally with mTFP fluorescence.

**Figure 5.**
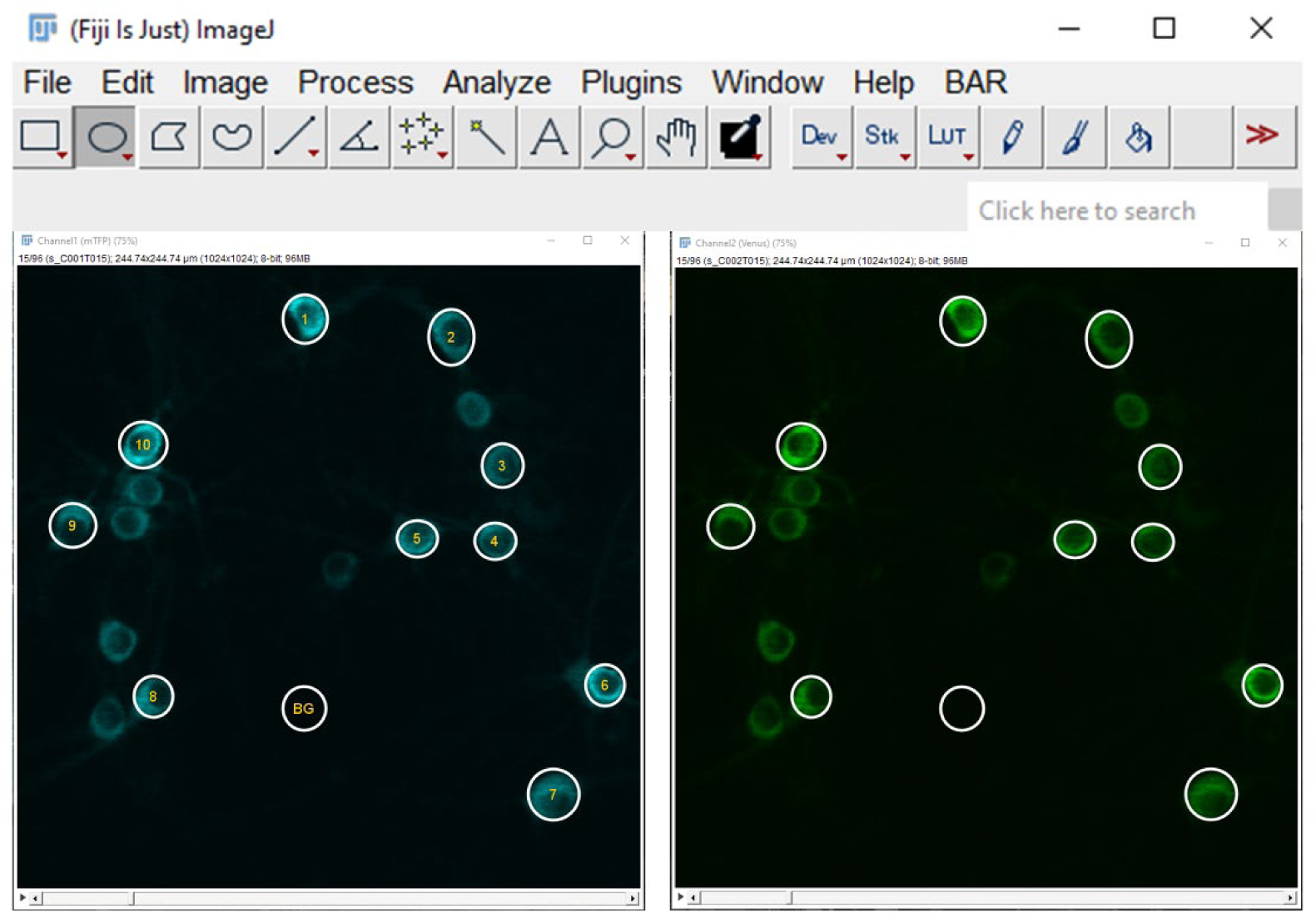
Screenshot of data acquisition for Pyronic FRET imaging using Fiji ImageJ. Cytosolic Pyronic mTFP and Venus signal from neurons. Regions of interest (ROIs) for analysis of specific individual cells (numbered circles) and background region between cells (BG circle) were selected. Background area is used to subtract background fluorescence from each signal.

**Figure 6.**
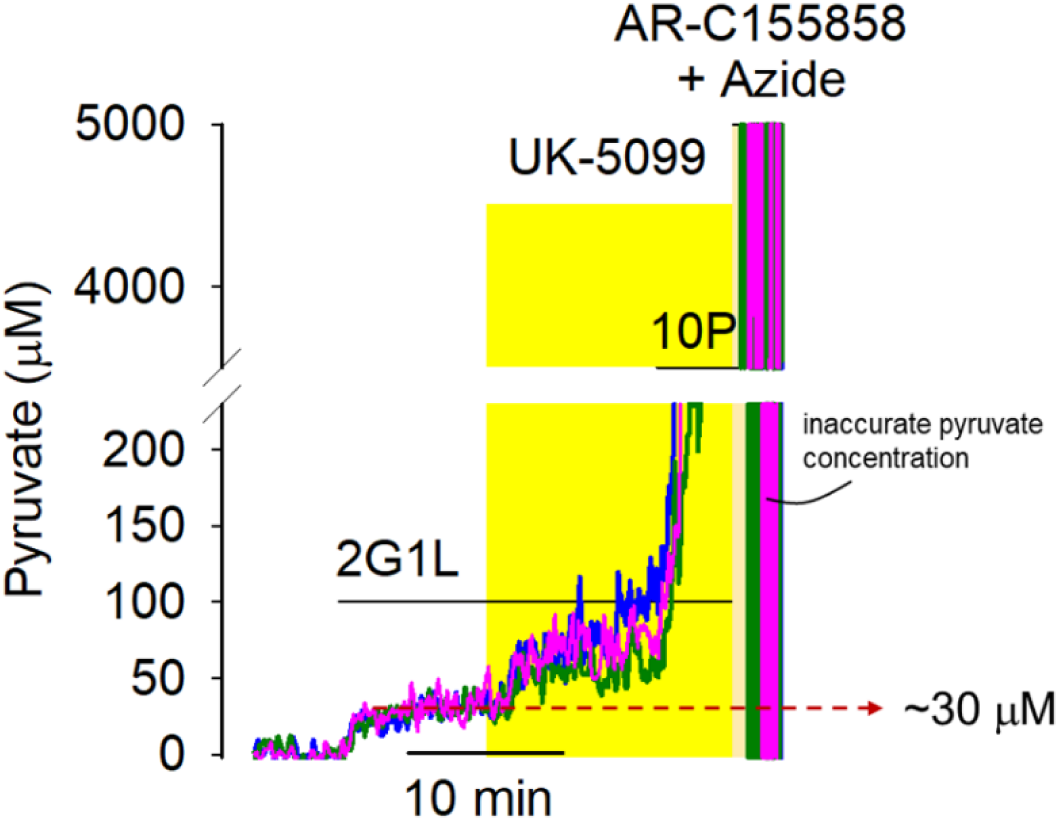
Calibrated signal of Pyronic. Cytosolic pyruvate concentration from three single hippocampal neurons obtained by the two-point calibration protocol. The reference buffer contains 2 mM glucose and 1 mM Lactate (2G1L).The original data that was used to calibrate the Pyronic signal is presented in figure 4B.

### 7.6 Pyruvate quantification

20. To compute the cytosolic pyruvate concentrations, the normalized Pyronic signal should be used to calculate the minimal FRET ratio R_MIN_ and the saturated FRET ratio R_MAX_. The *in vitro* Pyronic K_D_, which is 107 µM, should be obtained from the original paper of Pyronic [23]. These values should be included in the following equation

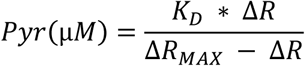

Where K_D_ corresponds to the dissociation constant obtained from an *in vitro* experiment with purified protein, ΔR is a given experimental point, and ΔR_MAX_ corresponds to the experimental dynamic range obtained from each cell. **Tip:** Given that the dynamic range (ΔR_MAX_) is determined for each experiment, FRET ratio values significantly close to the R_MAX_ will result in inaccurate pyruvate concentrations. This problem arises because the calibration equation’s denominator approaches zero when the Pyronic signal becomes saturated. However, physiological cytosolic pyruvate concentrations lie in the tens of micromolar to a few hundred micromolar under certain conditions, a safe range where the Pyronic signal is far from saturation.

## 8 RESULTS

### 8.1 Neuronal pyruvate concentration

Our two-point calibration method reveals that the Pyronic signal is approximately 4% above the minimum when neurons are bathed in glucose and lactate. This steady-state level likely results from the engagement of the glycolytic machinery and the backwards catalysis of LDH, which catabolized pyruvate in the cytosol. On the other hand, the pharmacological calibration raises the Pyronic signal to an approximate 20%, corresponding to the maximum Pyronic dynamic range when using lentivirus as a gene-delivery system. By using the in vitro K_D_ for Pyronic and the here-assessed ΔR_MAX_, the steady-state concentration of neuronal pyruvate is approximately 30 µM when neurons are supplied with 2 mM glucose and 1 mM lactate, which unveils a rather significant cytosolic pool of pyruvate in neurons.

## 9. LIMITATIONS

Certainly, two-point calibration has limitations. The introduced calibration protocol calculating cytosolic pyruvate concentration relies fundamentally on the assumed invariant K_D_. However, this parameter is derived from *in vitro* experiments using purified protein in an intracellular-mimicking buffer that approximates ionic strength but lacks the full complexity of the cellular environment. Consequently, possible discrepancies between the purified-protein and the *in cellulo* K_D_ should be taken into consideration. Nonetheless, potential different K_D_ will include a systematic error making the comparison in terms of fold-change secure to be analyzed.

The expression level of Pyronic might be a confounding factor, because it affects the indicator’s dynamic range. Therefore, a careful selection of MCT inhibitors to be used in the protocol is important, to correctly characterize the dynamic range of the fluorescent indicator in the cellular type of interest.

Although the absolute pyruvate concentration obtained from this protocol is informative, steady-state pyruvate concentration does not tell us too much about how cell metabolism is performing. Pyronic does not distinguish between glycolytic-derived and LDH-reversed pyruvate. Therefore, coupling this protocol with pharmacological perturbation of the steady-state enables a quantitative dissection of the fluxes that shape cellular pyruvate metabolism.

## 10. CONCLUSIONS

Pyruvate is a nodal intermediate in cellular metabolism. This protocol introduced a detailed procedure for quantifying cytosolic pyruvate concentration in neurons at single-cell resolution using a minimally invasive, two-point calibration approach employing the FRET-based genetically-encoded fluorescent indicator Pyronic. We envision that this protocol can be also used in other types of cells choosing the correct MCT inhibitor, allowing a quantitative assessment of pyruvate metabolism in mammalian systems. This facilitates comparisons across different cells, samples and experimental batches, thereby enabling comparison between a plethora of experimental conditions, which is not possible with intensity or FRET-ratio.

## Disclosure

The authors declare no conflicts of interest.

## Code - Data Availability

The datasets used and/or analyzed during the current study are available from the corresponding author upon reasonable request.

## Acknowledgments

This work was supported by an EMBO Postdoctoral Fellowship (EMBO ALTF 382-2021 to F. B.- L.). FONDECYT regular 1230145 and 1230682. ANID Doctoral Fellowships: 21200079 (Y.C) and 21190573 (C.A).

## Notes

### Competing Interest Statement

The authors have declared no competing interest.

## References

1. Acevedo A, Torres F, Kiwi M, Baeza-Lehnert F, Barros LF, Lee-Liu D, et al. Metabolic switch in the aging astrocyte supported via integrative approach comprising network and transcriptome analyses. Aging (Albany NY). 2023;15(19):9896–912. Epub 2023/04/19. doi: 10.18632/aging.204663. PubMed PMID: 37074814; PubMed Central PMCID: PMCPMC10599759.

2. Minhas PS, Jones JR, Latif-Hernandez A, Sugiura Y, Durairaj AS, Wang Q, et al. Restoring hippocampal glucose metabolism rescues cognition across Alzheimer’s disease pathologies. Science. 2024;385(6711):eabm6131. Epub 2024/08/22. doi: 10.1126/science.abm6131. PubMed PMID: 39172838.

3. Divakaruni AS, Wallace M, Buren C, Martyniuk K, Andreyev AY, Li E, et al. Inhibition of the mitochondrial pyruvate carrier protects from excitotoxic neuronal death. J Cell Biol. 2017;216(4):1091–105. Epub 2017/03/04. doi: 10.1083/jcb.201612067. PubMed PMID: 28254829; PubMed Central PMCID: PMCPMC5379957.

4. De La Rossa A, Laporte MH, Astori S, Marissal T, Montessuit S, Sheshadri P, et al. Paradoxical neuronal hyperexcitability in a mouse model of mitochondrial pyruvate import deficiency. Elife. 2022;11. Epub 2022/02/22. doi: 10.7554/eLife.72595. PubMed PMID: 35188099; PubMed Central PMCID: PMCPMC8860443.

5. Myeong J, Stunault MI, Klyachko VA, Ashrafi G. Metabolic regulation of single synaptic vesicle exo- and endocytosis in hippocampal synapses. Cell Rep. 2024;43(5):114218. Epub 2024/05/17. doi: 10.1016/j.celrep.2024.114218. PubMed PMID: 38758651; PubMed Central PMCID: PMCPMC11221188.

6. Isopi E, Granzotto A, Corona C, Bomba M, Ciavardelli D, Curcio M, et al. Pyruvate prevents the development of age-dependent cognitive deficits in a mouse model of Alzheimer’s disease without reducing amyloid and tau pathology. Neurobiol Dis. 2015;81:214–24. Epub 2014/12/02. doi: 10.1016/j.nbd.2014.11.013. PubMed PMID: 25434488.

7. Popova I, Malkov A, Ivanov AI, Samokhina E, Buldakova S, Gubkina O, et al. Metabolic correction by pyruvate halts acquired epilepsy in multiple rodent models. Neurobiol Dis. 2017;106:244–54. Epub 2017/07/16. doi: 10.1016/j.nbd.2017.07.012. PubMed PMID: 28709994.

8. Zilberter Y, Gubkina O, Ivanov AI. A unique array of neuroprotective effects of pyruvate in neuropathology. Front Neurosci. 2015;9:17. Epub 2015/03/06. doi: 10.3389/fnins.2015.00017. PubMed PMID: 25741230; PubMed Central PMCID: PMCPMC4330789.

9. San Martin A, Arce-Molina R, Aburto C, Baeza-Lehnert F, Barros LF, Contreras-Baeza Y, et al. Visualizing physiological parameters in cells and tissues using genetically encoded indicators for metabolites. Free Radic Biol Med. 2022;182:34–58. Epub 2022/02/21. doi: 10.1016/j.freeradbiomed.2022.02.012. PubMed PMID: 35183660.

10. Miyawaki A, Llopis J, Heim R, McCaffery JM, Adams JA, Ikura M, et al. Fluorescent indicators for Ca2+ based on green fluorescent proteins and calmodulin. Nature. 1997;388(6645):882–7. Epub 1997/08/28. doi: 10.1038/42264. PubMed PMID: 9278050.

11. Contreras-Baeza Y, Ceballo S, Arce-Molina R, Sandoval PY, Alegria K, Barros LF, et al. MitoToxy assay: A novel cell-based method for the assessment of metabolic toxicity in a multiwell plate format using a lactate FRET nanosensor, Laconic. PLoS One. 2019;14(10):e0224527. Epub 2019/11/02. doi: 10.1371/journal.pone.0224527. PubMed PMID: 31671132; PubMed Central PMCID: PMCPMC6822764.

12. Contreras-Baeza Y, Sandoval PY, Alarcon R, Galaz A, Cortes-Molina F, Alegria K, et al. Monocarboxylate transporter 4 (MCT4) is a high affinity transporter capable of exporting lactate in high-lactate microenvironments. J Biol Chem. 2019;294(52):20135–47. Epub 2019/11/14. doi: 10.1074/jbc.RA119.009093. PubMed PMID: 31719150; PubMed Central PMCID: PMCPMC6937558.

13. San Martin A, Ceballo S, Ruminot I, Lerchundi R, Frommer WB, Barros LF. A genetically encoded FRET lactate sensor and its use to detect the Warburg effect in single cancer cells. PLoS One. 2013;8(2):e57712. Epub 2013/03/08. doi: 10.1371/journal.pone.0057712. PubMed PMID: 23469056; PubMed Central PMCID: PMCPMC3582500.

14. Follenius-Wund A, Bourotte M, Schmitt M, Iyice F, Lami H, Bourguignon JJ, et al. Fluorescent derivatives of the GFP chromophore give a new insight into the GFP fluorescence process. Biophys J. 2003;85(3):1839–50. Epub 2003/08/29. doi: 10.1016/S0006-3495(03)74612-8. PubMed PMID: 12944297; PubMed Central PMCID: PMCPMC1303356.

15. Kremers GJ, Gilbert SG, Cranfill PJ, Davidson MW, Piston DW. Fluorescent proteins at a glance. J Cell Sci. 2011;124(Pt 2):157–60. Epub 2010/12/29. doi: 10.1242/jcs.072744. PubMed PMID: 21187342; PubMed Central PMCID: PMCPMC3037093.

16. Tsien RY. The green fluorescent protein. Annu Rev Biochem. 1998;67:509–44. Epub 1998/10/06. doi: 10.1146/annurev.biochem.67.1.509. PubMed PMID: 9759496.

17. Balleza E, Kim JM, Cluzel P. Systematic characterization of maturation time of fluorescent proteins in living cells. Nat Methods. 2018;15(1):47–51. Epub 2018/01/11. doi: 10.1038/nmeth.4509. PubMed PMID: 29320486; PubMed Central PMCID: PMCPMC5765880.

18. Liu B, Mavrova SN, van den Berg J, Kristensen SK, Mantovanelli L, Veenhoff LM, et al. Influence of Fluorescent Protein Maturation on FRET Measurements in Living Cells. ACS Sens. 2018;3(9):1735–42. Epub 2018/09/01. doi: 10.1021/acssensors.8b00473. PubMed PMID: 30168711; PubMed Central PMCID: PMCPMC6167724.

19. Waters JC. Accuracy and precision in quantitative fluorescence microscopy. J Cell Biol. 2009;185(7):1135–48. Epub 2009/07/01. doi: 10.1083/jcb.200903097. PubMed PMID: 19564400; PubMed Central PMCID: PMCPMC2712964.

20. Stepanenko OV, Stepanenko OV, Shcherbakova DM, Kuznetsova IM, Turoverov KK, Verkhusha VV. Modern fluorescent proteins: from chromophore formation to novel intracellular applications. Biotechniques. 2011;51(5):313–4, 6, 8 passim. Epub 2011/11/08. doi: 10.2144/000113765. PubMed PMID: 22054544; PubMed Central PMCID: PMCPMC4437206.

21. Lerchundi R, Huang N, Rose CR. Quantitative Imaging of Changes in Astrocytic and Neuronal Adenosine Triphosphate Using Two Different Variants of ATeam. Front Cell Neurosci. 2020;14:80. Epub 2020/05/07. doi: 10.3389/fncel.2020.00080. PubMed PMID: 32372916; PubMed Central PMCID: PMCPMC7186936.

22. Pathak D, Shields LY, Mendelsohn BA, Haddad D, Lin W, Gerencser AA, et al. The role of mitochondrially derived ATP in synaptic vesicle recycling. J Biol Chem. 2015;290(37):22325–36. Epub 2015/07/02. doi: 10.1074/jbc.M115.656405. PubMed PMID: 26126824; PubMed Central PMCID: PMCPMC4566209.

23. San Martin A, Ceballo S, Baeza-Lehnert F, Lerchundi R, Valdebenito R, Contreras-Baeza Y, et al. Imaging mitochondrial flux in single cells with a FRET sensor for pyruvate. PLoS One. 2014;9(1):e85780. Epub 2014/01/28. doi: 10.1371/journal.pone.0085780. PubMed PMID: 24465702; PubMed Central PMCID: PMCPMC3897509.

24. Comyn T, Preat T, Pavlowsky A, Placais PY. PKCdelta is an activator of neuronal mitochondrial metabolism that mediates the spacing effect on memory consolidation. Elife. 2024;13. Epub 2024/10/30. doi: 10.7554/eLife.92085. PubMed PMID: 39475218; PubMed Central PMCID: PMCPMC11524582.

25. Machler P, Wyss MT, Elsayed M, Stobart J, Gutierrez R, von Faber-Castell A, et al. In Vivo Evidence for a Lactate Gradient from Astrocytes to Neurons. Cell Metab. 2016;23(1):94–102. Epub 2015/12/25. doi: 10.1016/j.cmet.2015.10.010. PubMed PMID: 26698914.

26. Placais PY, de Tredern E, Scheunemann L, Trannoy S, Goguel V, Han KA, et al. Upregulated energy metabolism in the Drosophila mushroom body is the trigger for long-term memory. Nat Commun. 2017;8:15510. Epub 2017/06/06. doi: 10.1038/ncomms15510. PubMed PMID: 28580949; PubMed Central PMCID: PMCPMC5465319.

27. Rusu V, Hoch E, Mercader JM, Tenen DE, Gymrek M, Hartigan CR, et al. Type 2 Diabetes Variants Disrupt Function of SLC16A11 through Two Distinct Mechanisms. Cell. 2017;170(1):199–212 e20. Epub 2017/07/01. doi: 10.1016/j.cell.2017.06.011. PubMed PMID: 28666119; PubMed Central PMCID: PMCPMC5562285.

28. Baeza-Lehnert F, Saab AS, Gutierrez R, Larenas V, Diaz E, Horn M, et al. Non-Canonical Control of Neuronal Energy Status by the Na(+) Pump. Cell Metab. 2019;29(3):668–80 e4. Epub 2018/12/12. doi: 10.1016/j.cmet.2018.11.005. PubMed PMID: 30527744.

29. Wu D, Harrison DL, Szasz T, Yeh CF, Shentu TP, Meliton A, et al. Single-cell metabolic imaging reveals a SLC2A3-dependent glycolytic burst in motile endothelial cells. Nat Metab. 2021;3(5):714–27. Epub 2021/05/26. doi: 10.1038/s42255-021-00390-y. PubMed PMID: 34031595; PubMed Central PMCID: PMCPMC8362837.

30. Valdebenito R, Ruminot I, Garrido-Gerter P, Fernandez-Moncada I, Forero-Quintero L, Alegria K, et al. Targeting of astrocytic glucose metabolism by beta-hydroxybutyrate. J Cereb Blood Flow Metab. 2016;36(10):1813–22. Epub 2015/12/15. doi: 10.1177/0271678X15613955. PubMed PMID: 26661221; PubMed Central PMCID: PMCPMC5076786.

31. Lerchundi R, Fernandez-Moncada I, Contreras-Baeza Y, Sotelo-Hitschfeld T, Machler P, Wyss MT, et al. NH4(+) triggers the release of astrocytic lactate via mitochondrial pyruvate shunting. Proc Natl Acad Sci U S A. 2015;112(35):11090–5. Epub 2015/08/20. doi: 10.1073/pnas.1508259112. PubMed PMID: 26286989; PubMed Central PMCID: PMCPMC4568276.

32. San Martin A, Arce-Molina R, Galaz A, Perez-Guerra G, Barros LF. Nanomolar nitric oxide concentrations quickly and reversibly modulate astrocytic energy metabolism. J Biol Chem. 2017;292(22):9432–8. Epub 2017/03/28. doi: 10.1074/jbc.M117.777243. PubMed PMID: 28341740; PubMed Central PMCID: PMCPMC5454122.

33. Arce-Molina R, Cortes-Molina F, Sandoval PY, Galaz A, Alegria K, Schirmeier S, et al. A highly responsive pyruvate sensor reveals pathway-regulatory role of the mitochondrial pyruvate carrier MPC. Elife. 2020;9. Epub 2020/03/07. doi: 10.7554/eLife.53917. PubMed PMID: 32142409; PubMed Central PMCID: PMCPMC7077990.

34. Rabah Y, Frances R, Minatchy J, Guedon L, Desnous C, Placais PY, et al. Glycolysis-derived alanine from glia fuels neuronal mitochondria for memory in Drosophila. Nat Metab. 2023;5(11):2002–19. Epub 2023/11/07. doi: 10.1038/s42255-023-00910-y. PubMed PMID: 37932430; PubMed Central PMCID: PMCPMC10663161.

35. Ovens MJ, Davies AJ, Wilson MC, Murray CM, Halestrap AP. AR-C155858 is a potent inhibitor of monocarboxylate transporters MCT1 and MCT2 that binds to an intracellular site involving transmembrane helices 7-10. Biochem J. 2010;425(3):523–30. Epub 2009/11/26. doi: 10.1042/BJ20091515. PubMed PMID: 19929853; PubMed Central PMCID: PMCPMC2811425.

36. Polanski R, Hodgkinson CL, Fusi A, Nonaka D, Priest L, Kelly P, et al. Activity of the monocarboxylate transporter 1 inhibitor AZD3965 in small cell lung cancer. Clin Cancer Res. 2014;20(4):926–37. Epub 2013/11/28. doi: 10.1158/1078-0432.CCR-13-2270. PubMed PMID: 24277449; PubMed Central PMCID: PMCPMC3929348.

37. Halestrap AP. The mitochondrial pyruvate carrier. Kinetics and specificity for substrates and inhibitors. Biochem J. 1975;148(1):85–96. Epub 1975/04/01. doi: 10.1042/bj1480085. PubMed PMID: 1156402; PubMed Central PMCID: PMCPMC1165509.

38. Fernandez-Moncada I, Barros LF. Non-preferential fuelling of the Na(+)/K(+)-ATPase pump. Biochem J. 2014;460(3):353–61. Epub 2014/03/29. doi: 10.1042/BJ20140003. PubMed PMID: 24665934.

39. Barres BA, Koroshetz WJ, Chun LL, Corey DP. Ion channel expression by white matter glia: the type-1 astrocyte. Neuron. 1990;5(4):527–44. Epub 1990/10/01. doi: 10.1016/0896-6273(90)90091-s. PubMed PMID: 1698397.

40. Mamczur P, Borsuk B, Paszko J, Sas Z, Mozrzymas J, Wisniewski JR, et al. Astrocyte-neuron crosstalk regulates the expression and subcellular localization of carbohydrate metabolism enzymes. Glia. 2015;63(2):328–40. Epub 2014/09/27. doi: 10.1002/glia.22753. PubMed PMID: 25257920.

41. Looser ZJ, Faik Z, Ravotto L, Zanker HS, Jung RB, Werner HB, et al. Oligodendrocyte-axon metabolic coupling is mediated by extracellular K(+) and maintains axonal health. Nat Neurosci. 2024;27(3):433–48. Epub 2024/01/25. doi: 10.1038/s41593-023-01558-3. PubMed PMID: 38267524; PubMed Central PMCID: PMCPMC10917689.

42. Grimm D, Lee JS, Wang L, Desai T, Akache B, Storm TA, et al. In vitro and in vivo gene therapy vector evolution via multispecies interbreeding and retargeting of adeno-associated viruses. J Virol. 2008;82(12):5887–911. Epub 2008/04/11. doi: 10.1128/JVI.00254-08. PubMed PMID: 18400866; PubMed Central PMCID: PMCPMC2395137.

43. McMahon SM, Jackson MB. An Inconvenient Truth: Calcium Sensors Are Calcium Buffers. Trends Neurosci. 2018;41(12):880–4. Epub 2018/10/06. doi: 10.1016/j.tins.2018.09.005. PubMed PMID: 30287084; PubMed Central PMCID: PMCPMC6252283.

44. Akerboom J, Rivera JD, Guilbe MM, Malave EC, Hernandez HH, Tian L, et al. Crystal structures of the GCaMP calcium sensor reveal the mechanism of fluorescence signal change and aid rational design. J Biol Chem. 2009;284(10):6455–64. Epub 2008/12/23. doi: 10.1074/jbc.M807657200. PubMed PMID: 19098007; PubMed Central PMCID: PMCPMC2649101.

45. Gasterstadt I, Jack A, Stahlhut T, Rennau LM, Gonda S, Wahle P. Genetically Encoded Calcium Indicators Can Impair Dendrite Growth of Cortical Neurons. Front Cell Neurosci. 2020;14:570596. Epub 2020/11/17. doi: 10.3389/fncel.2020.570596. PubMed PMID: 33192315; PubMed Central PMCID: PMCPMC7606991.

46. Barakat S, Cimen S, Miri SM, Vatandaslar E, Yelkenci HE, San Martin A, et al. Bioenergetic shift and proteomic signature induced by lentiviral-transduction of GFP-based biosensors. Redox Biol. 2024;78:103416. Epub 2024/11/13. doi: 10.1016/j.redox.2024.103416. PubMed PMID: 39509993; PubMed Central PMCID: PMCPMC11574814.

47. Somero GN. Adaptation of enzymes to temperature: searching for basic “strategies”. Comp Biochem Physiol B Biochem Mol Biol. 2004;139(3):321–33. Epub 2004/11/17. doi: 10.1016/j.cbpc.2004.05.003. PubMed PMID: 15544958.

48. Bulusu V, Prior N, Snaebjornsson MT, Kuehne A, Sonnen KF, Kress J, et al. Spatiotemporal Analysis of a Glycolytic Activity Gradient Linked to Mouse Embryo Mesoderm Development. Dev Cell. 2017;40(4):331–41 e4. Epub 2017/03/02. doi: 10.1016/j.devcel.2017.01.015. PubMed PMID: 28245920; PubMed Central PMCID: PMCPMC5337618.

49. Harada K, Chihara T, Hayasaka Y, Mita M, Takizawa M, Ishida K, et al. Green fluorescent protein-based lactate and pyruvate indicators suitable for biochemical assays and live cell imaging. Sci Rep. 2020;10(1):19562. Epub 2020/11/13. doi: 10.1038/s41598-020-76440-4. PubMed PMID: 33177605; PubMed Central PMCID: PMCPMC7659002.

50. Imai S, Hario S, Suerte C, Yamaguchi I, Sakoi K, Terai T, et al. High-performance genetically-encoded green and red fluorescent biosensors for pyruvate. bioRxiv. 2025:2025.04.17.649293. doi: 10.1101/2025.04.17.649293.

51. Schindelin J, Arganda-Carreras I, Frise E, Kaynig V, Longair M, Pietzsch T, et al. Fiji: an open-source platform for biological-image analysis. Nat Methods. 2012;9(7):676–82. Epub 2012/06/30. doi: 10.1038/nmeth.2019. PubMed PMID: 22743772; PubMed Central PMCID: PMCPMC3855844.

52. Ai HW, Henderson JN, Remington SJ, Campbell RE. Directed evolution of a monomeric, bright and photostable version of Clavularia cyan fluorescent protein: structural characterization and applications in fluorescence imaging. Biochem J. 2006;400(3):531–40. Epub 2006/07/25. doi: 10.1042/BJ20060874. PubMed PMID: 16859491; PubMed Central PMCID: PMCPMC1698604.

53. Nagai T, Ibata K, Park ES, Kubota M, Mikoshiba K, Miyawaki A. A variant of yellow fluorescent protein with fast and efficient maturation for cell-biological applications. Nat Biotechnol. 2002;20(1):87–90. Epub 2001/12/26. doi: 10.1038/nbt0102-87. PubMed PMID: 11753368.

54. Day RN, Booker CF, Periasamy A. Characterization of an improved donor fluorescent protein for Forster resonance energy transfer microscopy. J Biomed Opt. 2008;13(3):031203. Epub 2008/07/08. doi: 10.1117/1.2939094. PubMed PMID: 18601527; PubMed Central PMCID: PMCPMC2483694.

